# Super-resolution going viral: T4 virus particles as perfect nature-designed 3D-Bio-NanoRulers

**DOI:** 10.1101/2024.04.04.588072

**Authors:** José Ignacio Gallea, Oleksii Nevskyi, Zuzanna Kaźmierczak, Tao Chen, Paulina Miernikiewicz, Anna Chizhik, Krystyna Dąbrowska, Mark Bates, Jörg Enderlein

## Abstract

In the burgeoning field of super-resolution fluorescence microscopy, significant efforts are being dedicated to expanding its applications into the three-dimensional domain. Various methodologies have been developed that enable isotropic resolution at the nanometer scale, facilitating the visualization of three-dimensional subcellular structures with unprecedented clarity. Central to this progress is the need for reliable 3D structures that are biologically compatible for validating resolution capabilities. Choosing the optimal standard poses a considerable challenge, necessitating, among other attributes, precisely defined geometry and the capability for specific labeling at sub-diffraction-limit distances.

In this context, we introduce the use of the non-human-infecting virus, bacteriophage T4, as an effective and straightforward bio-ruler for three-dimensional super-resolution imaging. Employing DNA point accumulation for imaging in nanoscale topography (DNA-PAINT) along with the technique of astigmatic imaging, we uncover the icosahedral capsid of the bacteriophage T4, measuring 120 nm in length and 86 nm in width, and its hollow viral tail. This level of detail in light microscopy represents a significant advancement. We further outline a simple protocol for the production and preparation of samples. Moreover, we explore the extensive potential of bacteriophage T4 as a multi-faceted 3D bio-ruler, proposing its application as a novel benchmark for three-dimensional super-resolution imaging in biological studies.

## 1. Introduction

The advent of super-resolution fluorescence microscopy techniques, breaking through the diffraction limit of light, has revolutionized our ability to visualize life at its most fundamental level. Among these, Single Molecule Localization Microscopy (SMLM) methods stand out for their capacity to delineate biological structures, such as cellular organelles and protein complexes, with near-molecular resolution.^[1–2]^ These techniques have illuminated the once obscure realm of ‘sub-microscopic’ infectious agents, namely viruses, which conventional light microscopy struggled to detail.^[3]^

Initially confined to two-dimensional imaging, these pioneering technologies have expanded into the three-dimensional (3D) sphere, enhancing our understanding of biological specimens’ true complexities.^[4–7]^ Recent advancements have achieved isotropic resolution within the nanometer range, enabling the precise visualization of 3D structures.^[8–11]^

Despite the widespread adoption of SMLM techniques in biological research, they remain complex technologies, particularly for 3D imaging. Consequently, tools and methods that validate the accuracy and reliability of these techniques are essential. Scientists have employed various strategies to tackle this, including image analysis and experimental methods. A common approach involves measuring localization precision, which indicates the uncertainty in determining individual molecules’ positions.^[12]^ However, this can overestimate resolution capabilities by not accounting for factors such as label density and sample drift. Other methods, like Fourier Ring Correlation (FRC)^[13]^ and decorrelation analysis,^[14]^ offer direct resolution estimates from super-resolved images but may miss local variations and depend heavily on proper image capture and processing.

To bridge these gaps, researchers have turned to reference structures, often called “standards” or “rulers”. These standards help relate estimated emitter positions to true positions,^[12]^ enhancing localisation accuracy and spatial resolution estimation. These structures ideally possess a well-defined geometric arrangement, have multiple reference points strategically placed at distances below the diffraction limit, allow for common tagging schemes, and are stable and highly reproducible. The use of these rulers allows for correlating estimated positions of emitters to their true position established by the reference points, thereby checking the localisation accuracy. They also facilitate the estimation of spatial resolution by analysing the distances between fluorophores positioned at known intervals, effectively acting as molecular rulers.

DNA origami structures, with their customizable shapes and precise fluorophore placement, exemplify such standards. These structures are long single-stranded DNAs folded by using numerous short DNA ‘staple’ strands to form a well-defined 2D or 3D structure.^[15]^ DNA origami provides exceptional versatility, enabling the accurate positioning of fluorophores at specific sites and desired length scales. However, it is important to acknowledge that these structures are predominantly suited for cell-free environments and encounter certain constraints with regard to the selection of labels,^[16]^ and their preparation can be challenging. Another notable example involves employing cellular structures like microtubules or the nuclear pore complex (NPC) as reference standards. Microtubules, in particular, are frequently utilized to evaluate spatial resolution through the measurement of their cross-sectional fluorescence profile. This method facilitates the determination of peak-to-peak distances between fluorophores positioned on opposing sides of the microtubule’s hollow cylindrical structure. Nonetheless, this technique is constrained by the inherent width of the microtubules (approximately 25 nm, excluding linkage error), is prone to selection bias, and is limited to applications within cellular contexts.^[17]^ On the other hand, NPCs have become a widely used reference standard.^[16]^ These cylindrical protein complexes, situated in the nuclear envelope, offer a significant resource for resolution assessment. For instance, labelling NUP96, a component protein of NPCs, enables distance measurements ranging from approximately 12 nm to 107 nm under optimal conditions. This allows for direct evaluation of whether the resolution exceeds these distances. However, the application of NPCs is confined to cellular contexts and may present implementation challenges if not previously optimized within the research group.

In an excellent 2023 study, Helmrich et al. Introduce PCNAs as a protein-based nanoruler capable of achieving sub-10 nm resolution.^[18]^ This homo-trimeric protein, essential for DNA replication and repair and measuring 8 nm, has undergone genetic code expansion to include three labelling sites strategically spaced at 6 nm intervals. This modification enables precise labelling through biorthogonal click chemistry. Moreover, PCNA demonstrates remarkable stability within cellular environments, offering significant benefits in scenarios where accurate stoichiometry is crucial. However, one of the major limitations of this nanoruler is its inability to measure in three dimensions, its limited versatility and the need for knowledge of genetic code expansion technology.

In this work, we introduce the exceptional capability of the non-pathogenic virus, bacteriophage T4 (T4), to act as a versatile 3D-Bio-NanoRuler, enhancing the use of super-resolution fluorescence microscopy. This naturally occurring DNA-protein complex, with its icosahedral capsid measuring 120 nm in length and 86 nm in width, paired with a cylindrical hollow viral tail, forms a unique 260 nm geometric rocket-shaped structure. By employing the DNA point accumulation for imaging in nanoscale topography (DNA-PAINT)^[19–20]^, in tandem with optical astigmatism,^[21]^ we have achieved detailed visualization of T4’s 3D structure with an unparalleled level of clarity using light microscopy. Additionally, we demonstrate T4’s utility in assessing microscope performance, where the phage offers an extremely rigid three-dimensional structure for calibrating microscopes with nanometer accuracy. Our streamlined preparation method not only yields a high quantity of the virus but also ensures that the majority of viruses are oriented perpendicular to the coverslip, making them ideal for use as 3D standards. Finally, we investigate their use as universal single-digit nanometer rulers, underlining their significant potential for advancing nanoscale precision within biological research.

## 2. Results

### 2.1. 3D-Bio-NanoRuler sample preparation and 3D imaging

T4 are double-stranded DNA viruses of the *Straboviridae* family within *Caudoviricetes*, that infect *Escherichia coli* (E. coli) bacteria. Morphologically, T4 represents myoviruses, having an icosahedral head and a long, contractile but non-flexible tail. These viruses are ubiquitous in nature and can be safely manipulated in laboratories operating at the lowest biosafety level (BSL-1). This safety profile contributes to the widespread use of T4 as a model organism for biological research and as a standard tool for assessing aerosol containment in cell sorters.^[22]^ T4 viruses are readily available on the market and can be replicated and purified using several established simple protocols, two of which we have used and are described in the Experimental section.

Another fascinating feature of T4 is its ability to orient itself perpendicularly to an albumin-coated substrate, a phenomenon that, to our knowledge, has not been previously documented. We discovered this through the use of an easy sample preparation protocol outlined in Figure 1 and detailed in the Experimental section. This protocol consists of five main preparation steps for imaging: 1- coating the coverslip with bovine serum albumin (BSA); 2- applying the solution containing the viruses to the substrate; 3- promoting the interaction of T4 with the substrate by vacuum drying; 4- gently hydrating and fixing the sample; 5- immunostaining.

**Figure 1.**
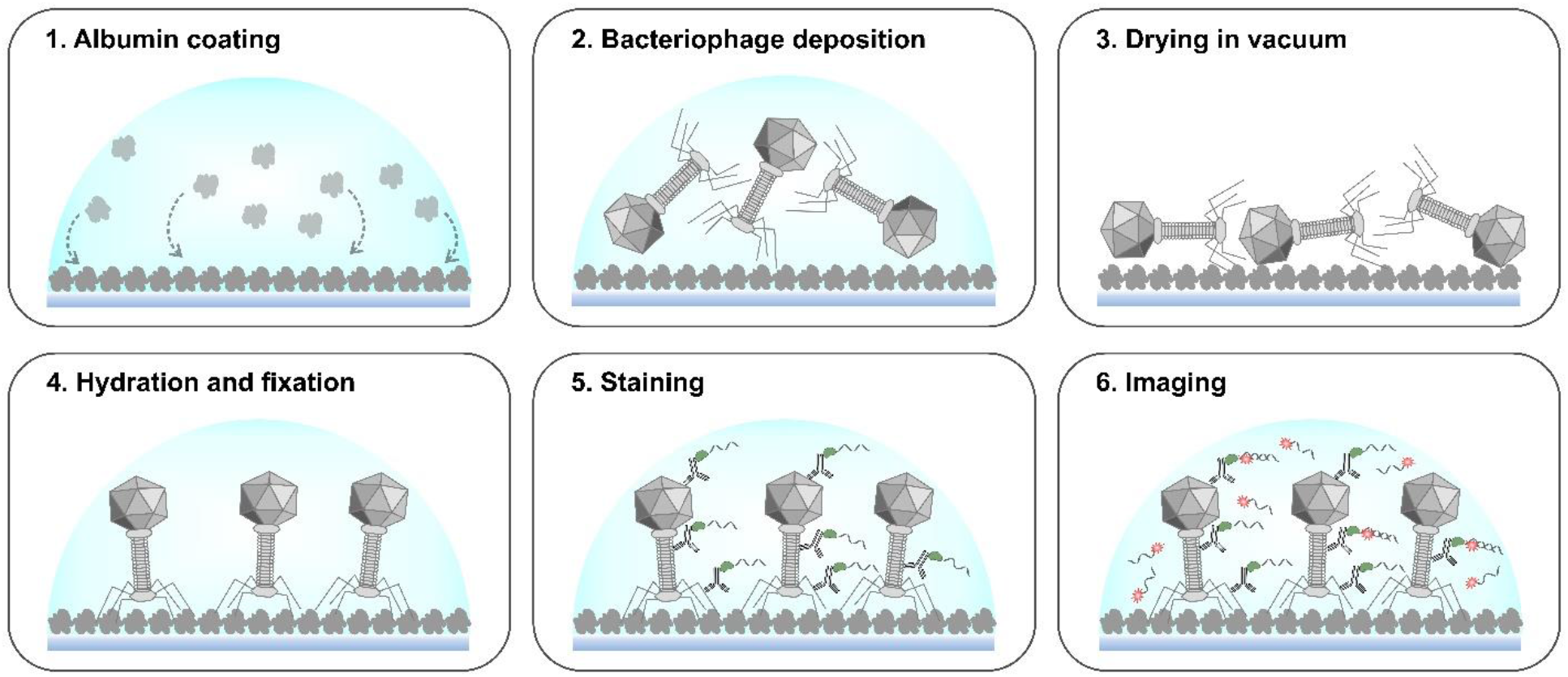
Preparation scheme of the T4 bacteriophage sample for DNA-PAINT imaging.

Following phage purification using the two different protocols, we confirmed their presence and quality by transmission electron microscopy (see Figure S1, Supporting Information) and assessed phage concentration. We used the phages purified by protocol 1 (see Experimental section) to prepare the sample according to our protocol, due to the higher purity observed under TEM. In this case, we labelled the phages with a solution containing T4-specific IgG antibodies precipitated from T4-specific serum of C57BL/6J mice (polyclonal) and secondary nanobodies conjugated to a DNA strand (Massive Photonics). We utilized a custom TIRF microscope to conduct 3D DNA-PAINT imaging, employing a solution containing a complementary DNA imager tagged with the Atto 643 fluorescent dye. To achieve the 3D modality, we integrate a cylindrical lens into the imaging setup, placing it along the microscope’s optical path.^[21]^ By analyzing the ellipticity and orientation changes of the fluorophore’s images it was possible to determine the z-coordinate of the fluorophore and thus reconstruct the corresponding 3D image. In Figure 2 and Video S1-2 we show, for the first time at this level of detail for light microscopy, the 3D architecture of T4. The characteristic elongated icosahedral hollow capsid and its cylindrical tail are clearly visible. We found that with this simple protocol hundreds of viruses were captured in sharp focus within a single field of view, with an exceptional signal-to-noise ratio. Strikingly, the majority of viruses (87%) were observed oriented perpendicular to the substrate, with their capsids clearly pointing upwards (Figure 2 and Video S1-3). The remaining 13% of viruses were observed with the capsids touching the substrate. By examining the TEM images of our samples, we discovered few cases where viruses appeared incomplete, lacking tails and fibers. This observation may explain the occurrence of some of these positions. There were a very small number of virus aggregates, and these were not included in the analysis.

**Figure 2.**
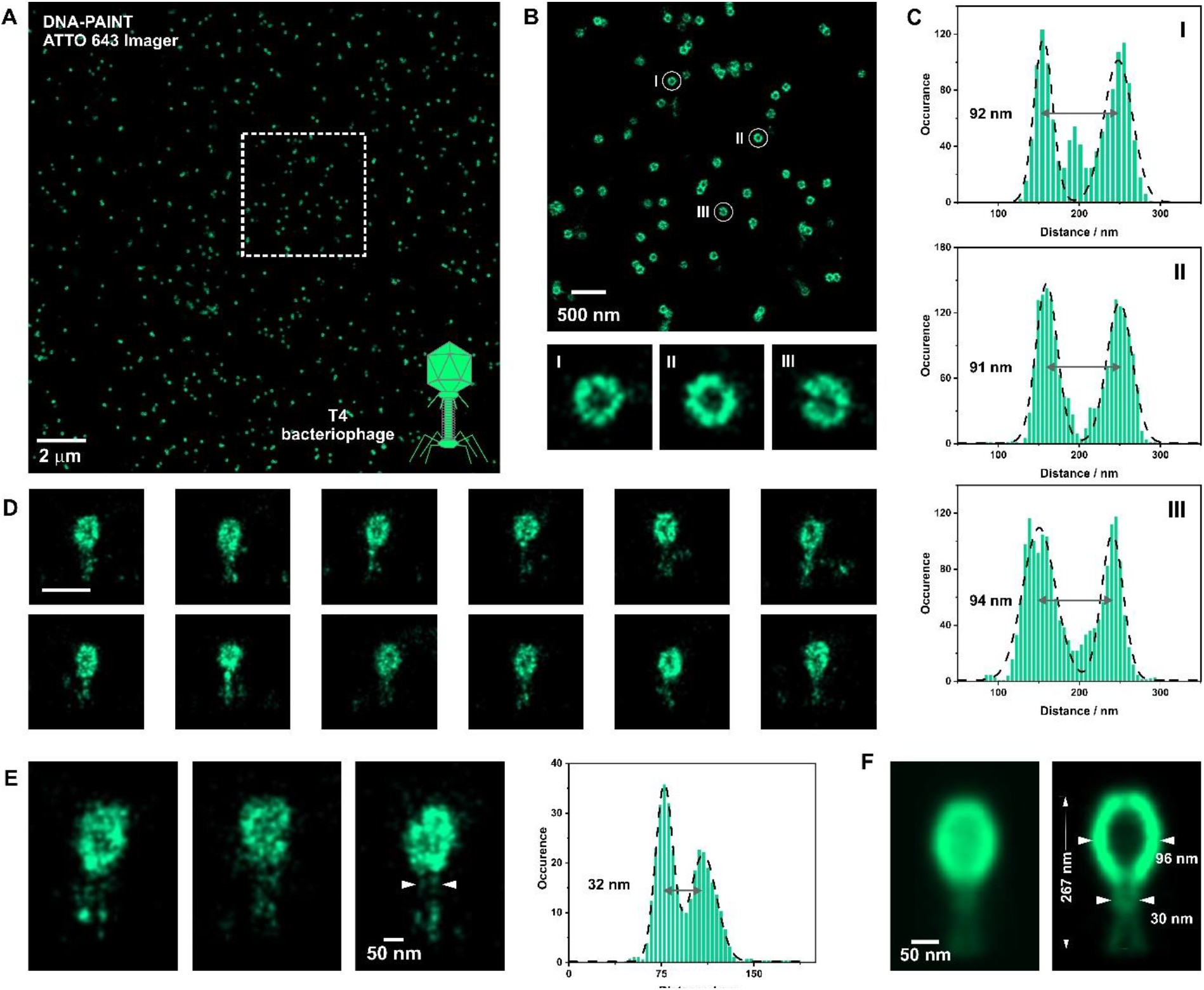
T4 bacteriophage 3D DNA-PAINT imaging. (a) Representative DNA-PAINT super-resolved image of the single viruses on the surface, where primary antibody recognized the whole phage. Imager was labeled with Atto 643 dye. (b) *x,y -* projection through the center of the capsid of the region marked in (a). Several individual phages I-III represented a hollow structure of the capsid. The scale bar is 100 nm. (c) Cross-sections through the center of the I-III capsids. (d) *xz*-projections of the individual phages represented in (a). The scale bar is 250 nm. (e) *xz*-projections of the individual phages where hollow structure of the virus tail could be observed. (f) Averaged *x,z*-projection of the phage dataset presented in (a).

We propose that the specific perpendicular orientation of the majority of T4 viruses, together with their particular known geometry and asymmetry in the horizontal plane (the upper part with respect to the lowest) and their versatility in handling, showcase some of the remarkable properties of this virus as a 3D-Bio-NanoRuler.

### 2.2. Resolution and image quality estimation with the 3D-Bio-NanoRuler

To validate the efficacy of our T4 virus as a 3D-Bio-NanoRuler, we performed an analysis of the capsid size in the lateral and axial dimensions. This included determining the resolution achieved by accurately measuring the distances between the capsid walls and comparing them to the actual size. This icosahedral structure, which contain the DNA of the virus, has rounded edges and measures 86 nm in width and 120 nm in length, as determined by cryo-EM.^[23]^ We generated 2D x-y representations from our 3D DNA-PAINT images, focusing on a specific range of z-positions that capture the widest span of the structure, particularly within the central region of the capsid (Figure 2 and Figure S3). Measuring the distances in the histograms of the head cross-sections, as in Figure 2C, yielded a width of 97±6 nm, which is consistent with the expected size considering a linkage error of ∼7 nm, as previously shown for the tandem primary antibody-secondary nanobody. Similarly, but in x-z representation, we verify the axial resolution by quantifying the width and also the length of the capsid and obtain values of ∼ 90 nm and ∼ 123 nm, respectively. These values were also consistent with the actual size of the virus head indicating that there were no depth-induced aberrations or an imperfect PSF calibration.

Another notable feature of T4 is its contractile tail, which can be precisely measured. When fully extended, this appendage is 140 nm long and 24 nm wide. In the region of the basal plate, close to the tail fibres or ‘legs’ of the virus, its width increases to 50 nm.^[24]^ It is noteworthy that in the majority of viruses observed, the hollow structure of the tail was discernible (Figure 2E), typically exhibiting an extended length of ∼140 nm and a width of ∼ 30 nm in the central region, while measuring ∼ 49 nm in the lower basal plate segment. Furthermore, these findings also aligned with the sizes obtained by measuring a virus particle resulting from merging the localization information of ∼650 viruses using a particle fusion approach (Figure 2F).^[25]^

Finally, we perform an FRC analysis on the super-resolved images and obtain an effective spatial resolution value of ∼3 nm (see an example in Figure S2). This indicates the good performance of our microscope and confirms that our 3D-Bio-NanoRuler measurements are within the resolution limits of our system.

### 2.3. Multi-target capability of T4

An inherent advantage of this naturally evolved structure is its vast display of targeting possibilities. T4 harbours over 40 unique structural proteins that form the head, neck, tail and fibres.^[24]^ Here, we use “Exchange 3D DNA-PAINT” to image first the entire virus and then a specific protein called fibritin. This protein, also known as gpwac, is a homotrimeric coiled-coil protein that forms six 53 nm long fibres or ‘whiskers’, that extend from the phage neck. This protein plays a key role in viral assembly and environmental sensing.^[26]^

In our two-step staining procedure, we began by labelling fibritin with a specific polyclonal rabbit anti-fibritin antibody and then labelled T4 with our previously used mouse polyclonal anti-T4 primary antibodies. This was done to minimize potential double labelling of the fibritin protein, as the mouse T4-specific IgG fraction may contain anti-fibritin antibodies. To complete the staining, we used a combination of secondary nanobodies: anti-mouse with an F1 DNA strand and anti-rabbit nanobodies with a different DNA strand (F2).

Our 2-colour xy plot shows colocalization of both labelled proteins. When analysing the 3D data, we observed the expected sub-capsid position of fibritin and a shape that resembles the fibrous collar of this protein (see Figure 3) with the expected size.

**Figure 3.**
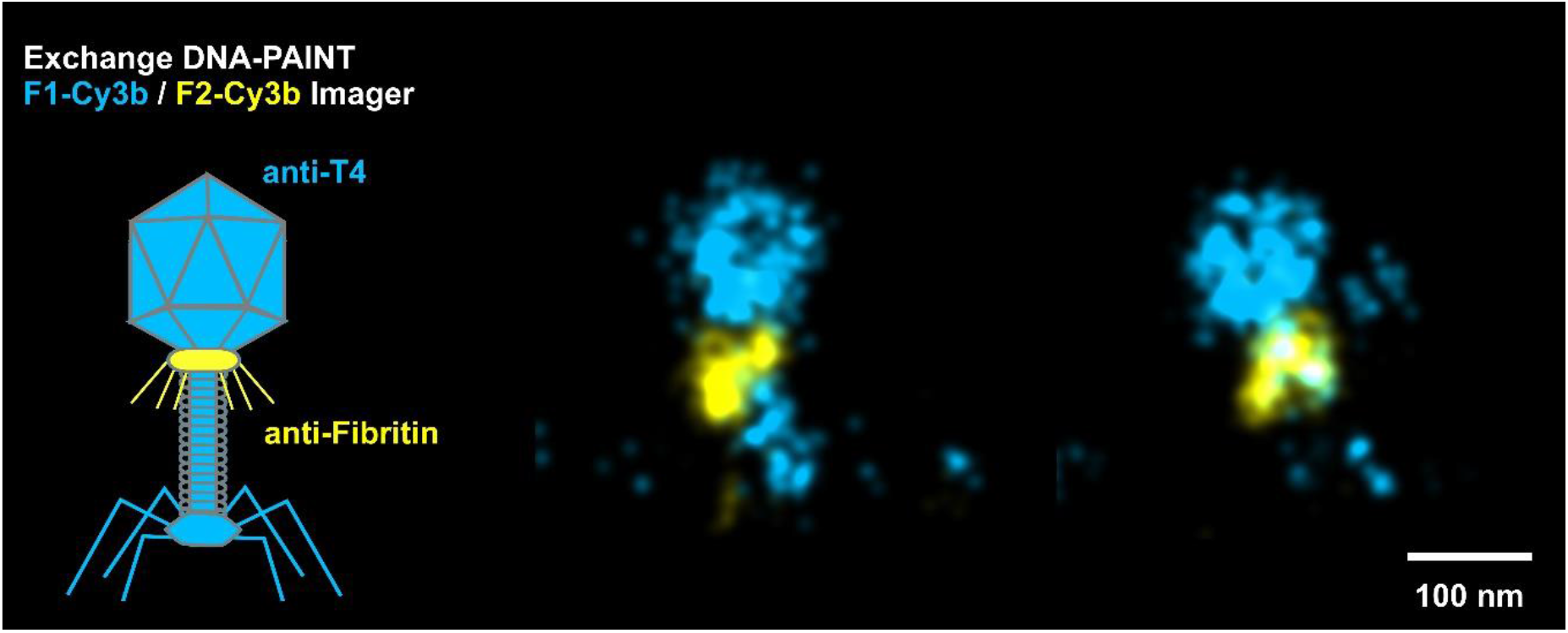
Dual color exchange DNA-PAINT imaging of the whole T4 bacteriophage (blue) and fibritin proteins (yellow).

### 2.3. Nanoruler versatility of T4 capsid

The remarkable and structurally stable T4 capsid offers a myriad of possibilities due to its unique geometry. Consisting of a 20-faced polyhedron with 10 equilateral end cap triangles and 10 scalene mid-section triangles, it is composed of four structural proteins: gp23, gp24, Hoc and Soc (Figure 4).^[23]^ Gp23, the most abundant protein, forms hexamers that constitute the surface lattice, while gp24, the head vertex protein, forms pentamers at eleven vertices (absent only at vertex where the tail is attached). Hoc is located at the center of each gp23 hexamer and Soc is distributed among gp23 with a hexagonal shape.

**Figure 4.**
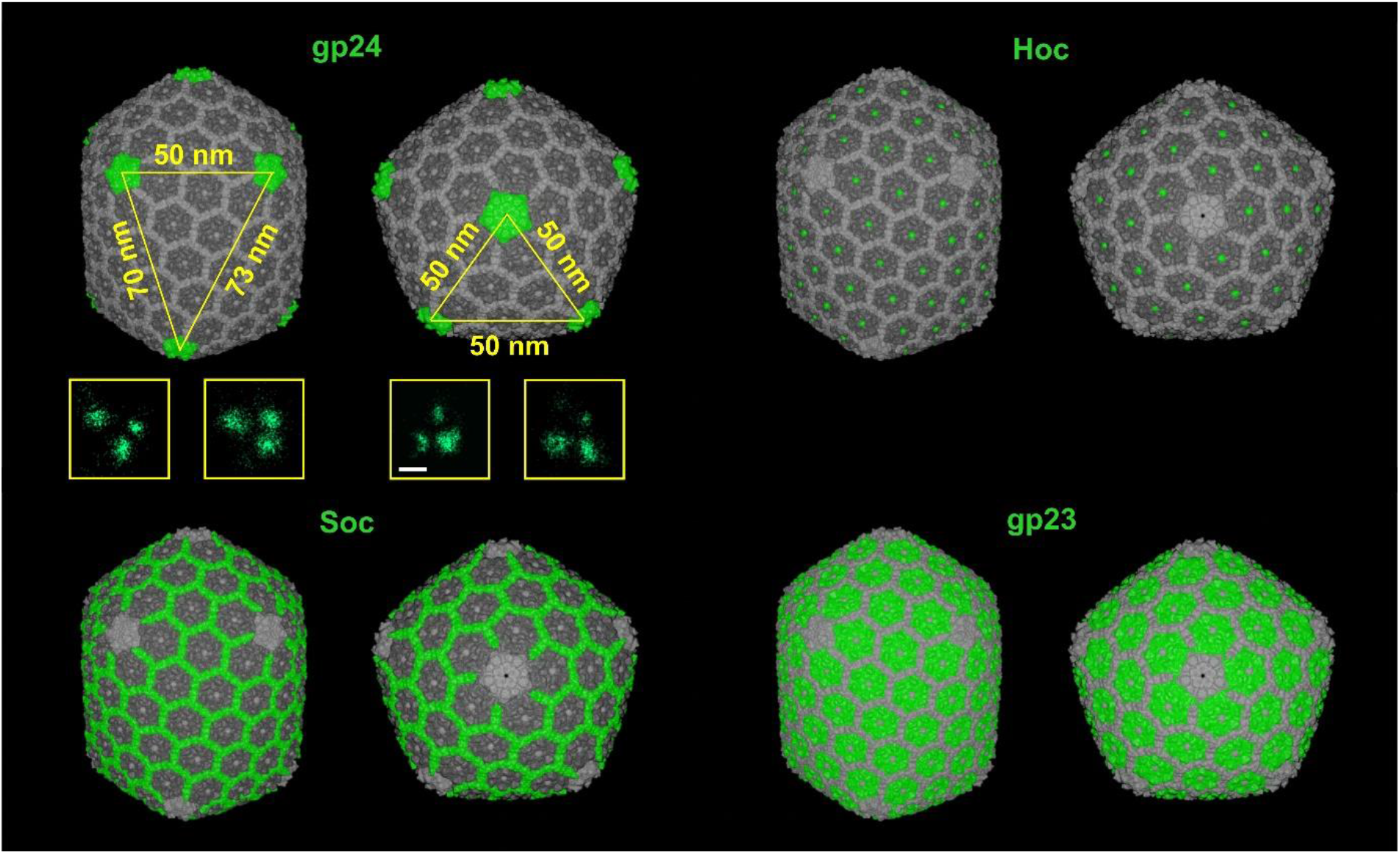
Side and top view of the T4 capsid is showing the 4 constituent proteins that can be labelled. Insets in yellow display the equilateral and scalene triangles obtained by DNA-PAINT imaging of gp24. The scale bar size is 50 nm. The 7VS5 pdb structure was used to generate this image.^[29]^

One avenue worth exploring is to use the immunolabeling of a protein such as gp24 to assess the effective labeling efficiency (ELE), in a similar way as done previously with the nuclear pore complex.^[16]^ The ELE indicates the proportion of target proteins that carry a fluorophore. This is a key factor as the accurate representation of biological samples in SMLM images is of paramount importance and is highly dependent on the density of fluorescent labelling.^[16]^ In addition, gp24 labelling may allow the measurement of smaller distances compared to the width of the capsid, facilitating quantification at higher resolutions. For example, measuring the distances between the corners of the scalene triangles (50, 70 and ∼73 nm) and the corners of the equilateral triangles (50 nm). In Figure 4 we show the two types of triangles generated through 2D DNA-PAINT imaging of the protein gp24. We measured 49 ± 3 nm for the equilateral triangle sides and, 53 ± 4 nm, 69 ± 3 and 74 ± 3 nm for the scaled triangle sides, indicating a good agreement with cryo-EM data.^[23]^

The T4 capsid also holds potential for measuring smaller distances, even down to the single-digit range. For example, it’s possible to label proteins such as Hoc, which maintain a separation of 14 nm between their counterparts,^[23]^ or Soc, which forms a hexagon of ∼14 nm from one side to the opposite. Another alternative is to label the six protrusions of the gp23 hexamer, which are 4.5 nm apart, or the five protrusions of the gp24 pentamer, which are 4.8 nm apart.^[23]^ In these cases, alternative labelling methods with minimal linkage error would be required. For example, immunostaining can be performed with primary nanobodies linked to a fluorescent probe or a DNA strand. Another interesting approach could be the expression of capsid proteins fused to self-labelling tags such as HaloTag or SNAP-tag. Notably, genetic modification of phages is well established in phage display technology, where exogenous peptides are fused to phage proteins, with T4 being one of the viruses used in this technology.^[27]^ In addition, the use of genetic code expansion for site-specific labelling, as demonstrated with the PCNA PicoRulers,^[18]^ is proving promising. Efficient site-specific incorporation of non-canonical amino acids into proteins has been achieved using M13 phages.^[28]^ All these advances pave the way for extending the 3D-Bio-NanoRuler capabilities of bacteriophage T4.

## 3. Conclusion/Discussion

The advent of super-resolution fluorescence microscopy has opened unprecedented opportunities in biological research while simultaneously presenting significant challenges. One of the primary hurdles is the need to make these advanced techniques easily and reliably accessible for widespread use. To address this, the development of standard structures, or “rulers,” has become an essential strategy for the smooth adoption of these innovative methods. These reference tools, ranging from specially designed protein structures and DNA constructs to cellular architectures, offer unique benefits and face distinct limitations. However, a universally adaptable ruler—simple to produce, utilize, flexible across various experimental contexts, and effective in three dimensions—has been conspicuously missing. In this study, we present the bacteriophage T4 as a versatile and universally applicable 3D-Bio-NanoRuler for super-resolution microscopy, showcasing its potential to navigate the complexities of microscopic measurement with ease and precision.

This virus, with its naturally designed protein nanostructure, is esteemed as a model organism in biological research. It has played a crucial role in enhancing our comprehension of life’s fundamental principles over the years. A notable instance of its contribution is the discovery of the triplet nature of the genetic code.^[30]^ This virus, which proliferates in Escherichia coli, thrives across a diverse array of environments inhabited by the bacterium, such as water bodies, soil, and even the gastrointestinal tracts of humans and animals. Recent findings have demonstrated that T4 can withstand varied temperature and pH levels, as well as proteolytic digestive enzymes, for at least 1-2 hours.^[31]^ This underscores T4’s remarkable resilience across a spectrum of environmental settings, encompassing physiological, non-physiological, and desiccated conditions.

We have devised a streamlined and user-friendly protocol for preparing 3D-Bio-NanoRulers for 3D SMLM imaging. Utilizing this protocol, we have attained an unprecedented level of detail in visualizing a virus using light microscopy. Previously, 3D imaging of Alexa Fluor 647 NHS Ester-labelled T7 phages through 4Pi single-molecule switching nanoscopy (W-4PiSMSN) was confined to visualizing only the virus’s head. Averaging techniques later enabled researchers to discern an icosahedral structure.^[32]^ On the other hand, 2D SMLM imaging of Alexa Fluor 647 NHS-labelled T4 facilitated a 3D phage reconstruction yet failed to reveal the hollow structures of the capsid and tail.^[33]^ In contrast, our 3D imaging technique allows for the direct observation of these distinctive hollow structures in T4 without the need for further post-processing or analysis.

A notable finding from our protocol is that a significant number of viruses display a perpendicular orientation to the substrate, mimicking the virus’s positioning during its infection process of *E. coli*. Although the precise dynamics of virus-host interaction remain under investigation, it is hypothesized that the interaction between the positively charged amino acids at the ends of the viral fibers and the negatively charged phosphate groups in the bacterial envelope’s lipopolysaccharides (LPS) plays a crucial role.^[34]^ The perpendicular stance of T4 in our samples may result from electrostatic attractions between the positively charged residues on the fibers and the negatively charged BSA used in the sample preparation. BSA carries a net negative charge at physiological pH levels^[35]^ and forms monolayers exhibiting a negative zeta potential.^[36]^

The unique orientation of T4 in our samples, previously undocumented, coupled with the virus’s notable stability under various conditions, underscores its potential as a reliable and versatile 3D ruler, even within cellular environments. Given the documented ability of phages to enter eukaryotic cells,^[37–38]^ there exists a promising avenue for employing T4 as an intracellular ruler. This application warrants further investigation and optimization.

In conclusion, the exceptional nanometer-scale resolution achievable in all dimensions and enhanced localization precision, along with its capacity for multi-target detection and single-digit nanometer accuracy, solidify T4’s position as a cutting-edge 3D-Bio-NanoRuler, showcasing unmatched versatility.

## 4. Experimental Section

### Bacteriophage

The T4 phage was acquired from the American Type Culture Collection (ATCC, Rockville, MD, USA) and cultured using Escherichia coli B obtained from the Collection of Microorganisms at the Institute of Immunology and Experimental Therapy, Wrocław, Poland, and from the Leibniz Institute DSMZ-German Collection of Microorganisms and Cell Cultures, Braunschweig, Germany, in LB-Broth high salt (Sigma-Aldrich). The culture was maintained at 37 °C for 8–10 hours. The phages were then purified with two different protocols: <u>Protocol 1</u>: Phage lysates were filtered through 0.22 µm polysulfone membrane filters (Merck Millipore, Billerica, MA, USA) and were used for subsequent purification and concentration by cross-flow filtration using Sartorius Hollow fiber cartridges. The phage preparation was further dialysed against PBS using 1000 kDa-pore membranes and purified by LPS-affinity chromatography EndoTrap Blue, following the manufacturer’s instructions (Hyglos GmbH, Bernried, Germany). LPS removal involved three successive incubations of the preparations with the slurry, followed by centrifugations. The final sample was dialyzed against PBS and filtered using 0.22 µm PVDF filters (Merck Millipore, Billerica, MA, USA). <u>Protocol 2</u>: The phage lysate was clarified by centrifugation at 3000 g for 20 min and incubated with 30 g/L NaCl and 75 g/L PEG 8000 at 4°C with continuous stirring. Phage particles were harvested by centrifugation at 3000g for 30 minutes at 4°C. The pellet was resuspended in PBS and the solution was mixed with an equal volume of chloroform. The solution was centrifuged at 5000g for 20 min and the aqueous upper phase was collected. The final sample was dialyzed against PBS and filtered using 0.22 µm PVDF filters (Merck Millipore, Billerica, MA, USA).

### Bacteriophage quantification

The purified phage preparation was assessed for phage concentration by determining the phage titer through serial dilution with PBS. Fifty microliters of each dilution were spotted on a culture plate pre-covered with susceptible bacteria, with three spots for each dilution. The plate was incubated at 37 °C for 8–10 hours until visible plaques appeared. The plaques were counted, mean values of three spots were calculated, and the phage concentration per milliliter was determined considering the dilution and spot volume.

### TEM imaging

A droplet of bacteriophage T4 solution (at a concentration of 10^12^ PFU/ml) was carefully absorbed onto carbon-coated copper grids (300 mesh, Ted Pella, Inc.), followed by a gentle wash with Milli-Q water. The grids were then stained with a 1% (w/v) uranyl acetate solution before imaging with a CM30 LaB6 transmission electron microscope (Philips).

### T4-specific serum production

C57BL/6J normal male mice (Mossakowski Medical Research Centre, Polish Academy of Sciences, Warsaw) were challenged with purified T4 phage to obtain T4 phage-specific serum. Three doses of highly purified phage preparation were administered intraperitoneally (IP) at 10^10^ pfu/mouse on days 0, 20 and 50. Blood samples were collected from the tail vein on day 100 and serum was separated as follows.

Serum was collected in tubes (BD SST II Advance), allowed to clot for 1 hour at room temperature (RT) and then separated from the clot by centrifugation (10 min, 2000 g). The separated serum was stored at −20°C until further use. The presence of T4-specific IgG antibodies in the serum was confirmed by ELISA.

### gp24 protein and its serum-specific production

The phage protein gp24 was produced as described by Miernikiewicz et al.^[39]^ Briefly, the gene encoding the mature form of the head vertex protein: gp24 was cloned into pDEST15 using Gateway technology, allowing expression of recombinant gp24 with a glutathione S-transferase (GST) affinity tag. gp24 was expressed in the Escherichia coli B834 (DE3) expression system. Expression of the recombinant phage protein was induced with 0.2 mM isopropyl-β-d-1-thiogalactopyranoside (IPTG) and carried out overnight at 25°C. Harvested bacteria were lysed with lysozyme in phosphate buffer containing phenylmethylsulfonyl fluoride (PMSF) (50 mM Na2HPO4, 300 mM NaCl, 1 mM PMSF; pH 7.5) and freeze-thawed. The soluble fraction of gp24 was incubated with glutathione sorbent slurry (Glutathione Sepharose 4B; GE Healthcare Life Sciences), washed with phosphate buffer, and phage protein was released by proteolysis with AcTev protease (5 U/ml) (Invitrogen, Life Technologies Corporation) at 10°C. Lipopolysaccharide (LPS) was then removed using EndoTrap HD (Lionex GmbH, Germany) and gel filtration fast protein liquid chromatography (FPLC) was performed on a Superdex 75 10/300 GL column (GE Healthcare Life Sciences). The sample was dialyzed against phosphate-buffered saline (PBS) and filtered through 0.22-µm polyvinylidene difluoride (PVDF) filters (Millipore). Protein concentration was determined by the Lowry chromogenic method (Fermentas International Inc.).

To induce anti-gp24 antibodies in mice (polyclonal), male C57BL/6J mice (Mossakowski Medical Research Centre, Polish Academy of Sciences, Warsaw) were challenged subcutaneously with three doses of the previously purified gp24 protein, 200 µg/mouse, on days 0, 20, and 40. Blood was collected from the orbital vein up to 7 days after the last dose.

Serum was collected in tubes (BD SST II Advance), allowed to clot for 1 h at RT, separated from the clot by centrifugation (10 min, 2000 g) and stored at −20°C until further use. The presence of gp24-specific IgG antibodies in the serum was confirmed by ELISA.

### Ammonium sulfate antibody precipitation

T4-specific and gp24-specific serum samples were incubated in a 50% ammonium sulfate saturated solution for 30 minutes on ice. The suspension was then centrifuged at 10,000g for 12 minutes at 2°C. The supernatant was removed, and the pellet was resuspended in PBS and stored at −20°C until further use.

### T4 sample preparation

Cleaned 18 mm coverslips (Paul Marienfeld, 0117580) were coated with 5% BSA solution in PBS (Blocking solution) for 2 hours at RT or overnight at 4°C, followed by three washes with PBS. Subsequently, 10 µl of T4 in PBS (8×10^11^ PFU/ml) was applied to the wet slides. The coverslips were then vacuum dried for 10 min in a centrifuge concentrator (Concentrator plus, Eppendorf) without rotation. The samples were then simultaneously hydrated and fixed with a solution of 4% PFA in PBS for 20 min, or alternatively gently hydrated with PBS for 5 min followed by the addition of 16% PFA solution in PBS to a final concentration of 4% PFA and incubated for 20 min. An optional quenching step was performed with 0.1 M glycine in PBS for 10 minutes. Finally, the coverslips were washed three times with PBS and blocked with Blocking solution for 1 hour at RT.

### T4 bacteriophage immunolabeling

After blocking, T4 samples were incubated with a 1:30 solution of mouse T4 specific IgG antibody in blocking solution for 1 hour for single target labeling. For dual target labeling, primary staining was performed in two steps: T4 samples were first incubated with a 1:50 solution of rabbit anti-wac (fibritin) antibody (2.95 mg/ml; CUSABIO, No. CSB-PA319157ZA01EDZ) in blocking solution for 1 hour, followed by incubation with a 1:30 solution of mouse T4 specific IgG antibody in blocking solution for 1 hour. The samples were washed three times for 10 minutes each with 0.1% Tween in PBS and then incubated for 45 minutes with a mixture of 1:50 DNA strand conjugated single domain secondary antibodies (FluoTag®-XM-QC anti-mouse IgG kappa light chain + FAST docking site F1 and FluoTag®-XM-QC anti-rabbit IgG + FAST docking site F2; Massive Photonics) in a solution of 5% BSA, 0.1% Triton-X in PBS. The samples were then washed three times for 10 minutes each with 0.1% Tween in PBS before imaging. An optional post-fixation step with 4% PFA (Sigma-Aldrich, F8775) in PBS for 10 minutes was performed in cases of long-term sample storage.

### DNA-PAINT imaging

2D DNA-PAINT imaging was performed on a custom built TIRF setup utilizing a 640 nm diode laser (Omicron Quixx, 140 mW). The laser beam was focused with a lens onto the back focal plane of a 100×/1.45 NA oil immersion objective (UPLXAPO100XO, Olympus). The lens in front of the microscope was mounted onto a micrometer translation stage (Thorlabs) to enable TIRF illumination of the sample by moving the laser beam parallel to the microscope. A custom-built three-axis linear stage was used for smooth lateral and axial sample positioning.

Additionally, objective was equipped with focus scanner (P-725.xCDE2 PIFOC, Physik Instrumente). The fluorescence was collected using the same objective and was spectrally separated from the excitation laser light by a quad-line beamsplitter (zt405/488/561/640rpc, AHF Analysentechnik Tübingen), further magnified by two lenses (*f*_1_ = 200 mm AC254-200-A and *f*_2_ = 400 mm AC254-400-A, both Thorlabs) and imaged onto the chip of an electron multiplying charge-coupled device (EMCCD) camera (Andor iXON Ultra 897). To suppress background stemming from the excitation laser, a notch filter (zet635NF, AHF Analysentechnik) and a bandpass filter (Brightline HC 708/75, AHF Analysentechnik) were placed between the two lenses, and a cleanup filter (Brightline HC 640/10, AHF Analysentechnik) was placed in front of the laser. The total magnification was 200-fold, resulting in an image with 70 nm per pixel of the CCD camera. All measurements were performed at 294 K. The obtained movies were analyzed to reconstruct super-resolution images with the Picasso software package.^[40]^

3D DNA-PAINT imaging was performed on the Olympus IX-71 based on inverted microscope stand and equipped with a UPLANSAPO 100X NA 1.4 oil-immersion objective lens. Laser illumination sources used for SMLM imaging included red and green lasers for imaging (642nm CW, 1.5W, MPB Communications Inc.; 532nm CW, 1.2W, Light Cube LC-LS-532-1.2W). Excitation light was controlled and modulated via an acousto-optic tunable filter (Crystal Technologies, AOTF-PCAOM). Variable angle TIRF or near-TIRF illumination was achieved using a custom light path entering through the rear port of the microscope. Excitation light was separated from fluorescence using a dichroic beam-splitter in the filter cube turret (650DCXR or Z532RDC, Chroma Technology Corp.) and an emission filter in the detection path (ET700/75 or HQ582/75, Chroma). Fluorescence light was collected by the objective lens, passed through an optical relay, and focused to form an image on a back-illuminated EMCCD camera (Andor Ixon+, DU860). For 3D imaging, a cylindrical lens (f = 500 mm) was placed in the optical path before the image plane. The microscope was equipped with a motorized sample stage (Märzhäuser Wetzlar), and the objective lens was mounted on a piezoelectric objective positioner (Piezo Jena). During image acquisition, the objective Z position was continuously adjusted to maintain a constant focus position. This focus-lock system was based on an infra-red laser beam introduced into the microscope via the right-side port below the filter turret and combined into the optical path using a short-pass dichroic mirror (900DCSP, Chroma). The focus lock laser (980nm, Thorlabs) was aligned to focus at the back focal plane of the objective lens and reflect from the glass-water interface of the sample. The position of the reflected beam was detected using a quandrant photodiode (Silicon Sensor Intl., QP50) which was monitored via a DAQ card (National Instruments). All microscope control and data acquisition was performed using custom software written in Labview (National Instruments). The sample was illuminated with 642nm or 532nm excitation light. The emitted light was filtered spectrally (see above) and detected at the EMCCD camera, running at a frame rate of 50 Hz. Typically, around 100,000 image frames were acquired in a single measurement. Optical stabilization of the z-focus (focus-lock) was engaged before starting each recording, in order to minimize sample drift during the measurement. Prior to the measurements, images of a fluorescent bead located on the sample were recorded as the bead was scanned in the Z-dimension, in order to create a calibration scan which was used in post-processing analysis of the image data.

For the single-target scenario, imaging was performed for 2 hours using the imaging buffer from the MASSIVE-SDAB-FAST 2-PLEX kit (Massive Photonics) containing 250 pM Atto 643-labelled Imager F1 supplemented with 80 nm gold nanoparticles (BBI Solutions) at a 1:10 dilution for drift correction. Exchange-DNA-PAINT strategy was utilized for the two-target scenario. First, the imaging buffer containing 250 pM Cy3b-labelled Imager F1 together with the gold nanoparticles was added and imaged for 2 hours. The sample was then carefully washed 2 times with 500 mM NaCl PBS (pH 8.0) and the imaging buffer containing 250 pM Cy3b-labelled Imager F2 was added and imaged for a further 2 hours.

### Data analysis

2D and 3D SMLM data were analyzed using custom software packages written by Dr. Mark Bates (unpublished). In general, SMLM image analysis and reconstruction follows a standard approach based on peak finding and localization. Correction of sample drift in post-processing was done based on image correlation of the SMLM data with itself over multiple time windows, using the COMET method (Reinkensmeier et al., manuscript in preparation). SMLM images were rendered as summed Gaussian spots (2D imaging) or as localization histograms (3D imaging) with a bin size typically chosen to be 6 nm. FRC analysis was done utilizing FIRE plugin for ImageJ. 3D data visualization (see Supplementary Information) was performed utilizing ViSP. FRC analysis was done utilizing FIRE plugin for ImageJ. 3D data video (see Supplementary Information) was performed utilizing ViSP.^[41]^

### Ethics Statements

All animal experiments were performed according to EU Directive 2010/63/EU for animal experimentations and were approved by the 1st Local Committee for Experiments with the Use of Laboratory Animals, Wroclaw, Poland (No. 64/2009 and 76/2011). The authors followed the ARRIVE (Animal Research: Reporting of in vivo Experiments) guidelines.^[42]^

## Acknowledgements

JIG acknowledges financial support from the European Union’s Horizon 2021 research and innovation program under the Marie Skłodowska-Curie Grant Agreement no.101062508 (project name: SOADOPP). JE and JIG acknowledge financial support by the DFG through Germany’s Excellence Strategy EXC 2067/1-390729940. JE and ON thank the European Research Council (ERC) for financial support via project “smMIET” (grant agreement no. 884488) under the European Union’s Horizon 2020 research and innovation program. We would like to thank Dr. Jan Christoph Thiele for the stimulating discussions at the beginning of the project. Special thanks also go to Dr. Alexey Chizhik for the creation of Figure 4.

## Conflict of Interest

The authors declare no conflict of interest.

## Data Availability Statement

The data that support the findings of this study are available from the corresponding author upon reasonable request.

## Notes

### Competing Interest Statement

The authors have declared no competing interest.

