## Supporting Information for "Super-resolution going viral: T4 virus particles as perfect nature-designed 3D-Bio-NanoRulers"

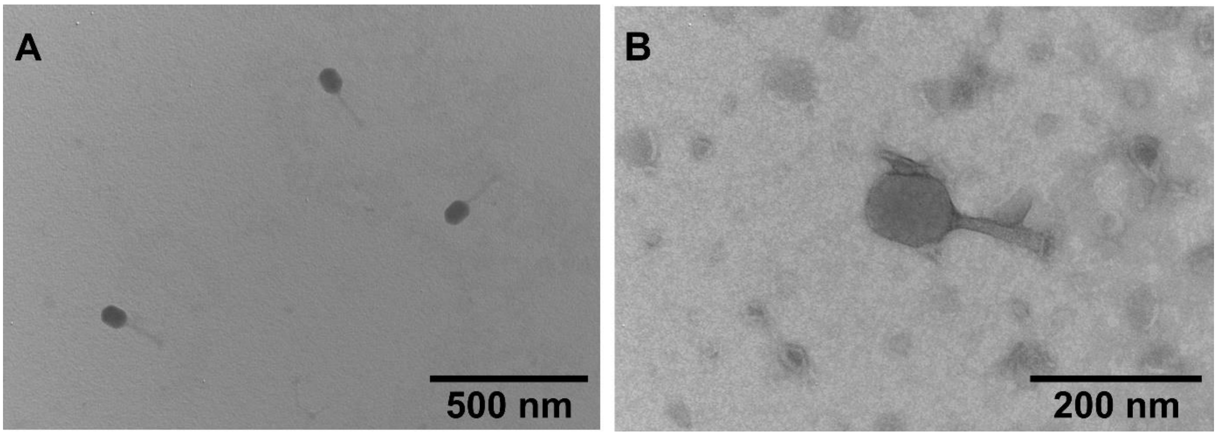


**Figure S1:** TEM images of T4 viruses purified by protocol 1 (A) and protocol 2 (B).

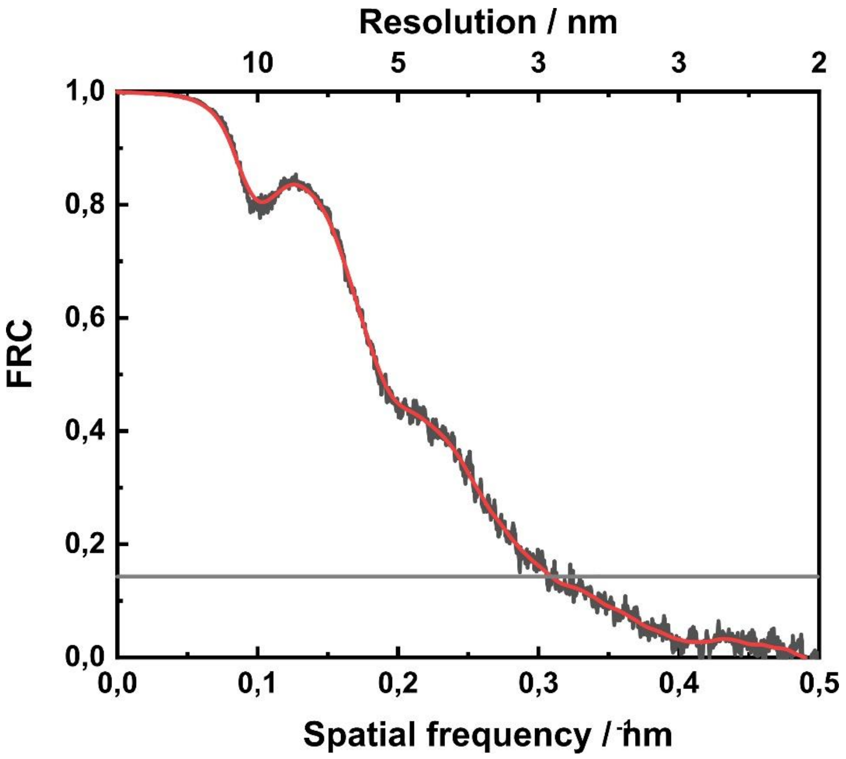


**Figure S2:** Fourier ring correlation analysis of super-resolved image presented in Figure 2.

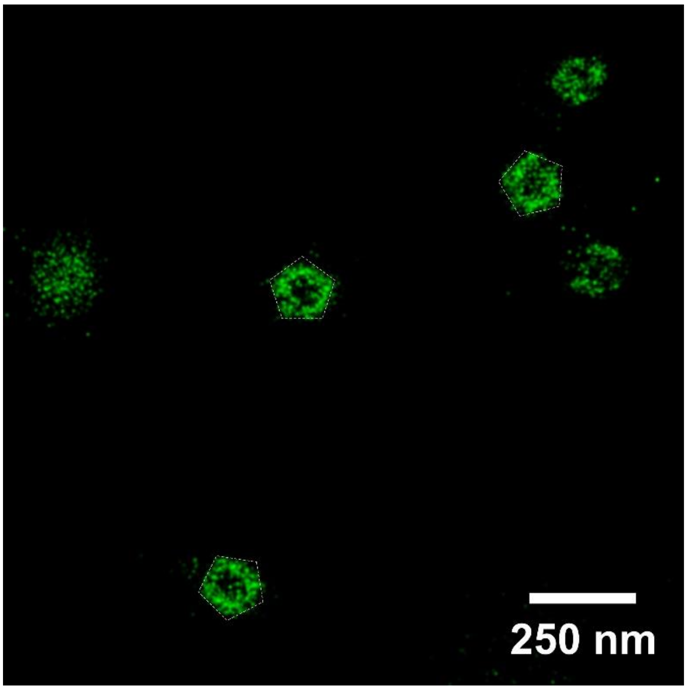


**Figure S3:** xy Cross-section of the phage head showing the hollow structure and its pentagonal shape (white dashed line).

**Video S1-S2:** T4 particles 3D representation.
